# Mitotic chromatin marking governs asymmetric segregation of DNA damage

**DOI:** 10.1101/2023.09.04.556166

**Authors:** Juliette Ferrand, Juliette Dabin, Odile Chevallier, Matteo Kane-Charvin, Ariana Kupai, Joel Hrit, Scott B. Rothbart, Sophie E. Polo

## Abstract

The faithful segregation of intact genetic material and the perpetuation of chromatin states through mitotic cell divisions are pivotal for maintaining cell function and identity across cell generations. However, most exogenous mutagens generate long-lasting DNA lesions that are segregated during mitosis. How this segregation is controlled is unknown. Here, we uncover a mitotic chromatin-marking pathway that governs the segregation of UV-induced damage in human cells. Our mechanistic analyses reveal two layers of control: histone ADP-ribosylation, and the incorporation of newly synthesized histones at UV damage sites, that both prevent local mitotic phosphorylations on histone H3 serine residues. Functionally, this chromatin-marking pathway drives the asymmetric segregation of UV damage in the cell progeny with consequences on daughter cell fate. We propose that this mechanism may help preserve the integrity of stem cell compartments during asymmetric cell divisions.

## INTRODUCTION

Genome integrity is constantly challenged by external and internal sources of genotoxic stress^1,2^. Cells have evolved elaborate pathways to repair DNA damage and to monitor genome integrity at critical steps of the cell cycle, thus preventing the transmission of damaged material across cell divisions^3–5^. Nevertheless, some DNA lesions escape classical checkpoint control and are transmitted to the cell progeny, sometimes unevenly, with adverse consequences on daughter cells inheriting the damage^6^. In particular, long-lasting DNA damage generated by exogenous mutagens such as UV light persist through mitosis, and their non-random segregation through multiple cell generations shapes cancer genome evolution^7^. Replication-associated endogenous damage, including under-replicated DNA, can also be passed on to the next cell cycle to be resolved in daughter cells^6,8,9^. Non-random segregation of replicative damage is dependent on the ATR-Chk1 kinase pathway^6,10^. However, control mechanisms for exogenous DNA damage segregation during cell division are unknown. The DNA damage response is accompanied by a reshaping of the chromatin landscape, including critical changes in histone post-translational modifications (PTMs)^11^. It is thus tempting to speculate that alterations in histone PTMs closely associated with mitotic events^12^ may contribute to control the segregation of DNA damage during cell division. Supporting this hypothesis, the mitotic histone mark threonine 3 phosphorylation on histone H3 (H3T3ph) drives asymmetric segregation of chromatin during the division of Drosophila male germline stem cells^13^. We thus decided to examine H3 mitotic phosphorylations during the response of human cells to UV-induced damage and their potential consequences on DNA damage segregation.

## RESULTS

### Non-random segregation of UV-damaged chromosomes through mitosis

Cells locally damaged with UV still divide and transmit the damaged material to the progeny, resulting in an easily identifiable scattered pattern of DNA damage in daughter cells (Fig.1a, red dots in G1 cell nuclei). To evaluate if post-replicative UV lesions are transmitted symmetrically or asymmetrically between daughter cells, we inflicted local UVC irradiation to cells synchronized at the G2/M boundary, which generated 1 to 4 UV damage spots per nucleus (Extended Data Fig.S1a), released the cells through mitosis and measured the levels of the main UV photoproducts, CPDs (Cyclobutane Pyrimidine Dimers), in early G1 cells (Fig. 1a) after CPD staining with a highly specific antibody (Extended Data Fig. S1b). Of note, the G1 daughter cell pairs were easily identified based on their spatial proximity, which we further validated based on microtubule staining (Extended Data Fig. S1c), and CPD signals show a linear correlation with the UV dose and thus are reliable measurements of UV-induced DNA damage (Extended Data Fig.S1d). We calculated the CPD bias in daughter cell pairs, which reflects the degree of asymmetry in UV damage distribution between daughter cells. We observed a wide range of UV damage segregation modes, from almost completely symmetric (ca. 1% CPD bias), meaning that daughter cells inherit similar amounts of DNA damage, to very asymmetric (ca. 60% CPD bias), where most of the DNA damage ends up in one daughter cell (Fig.1a). CPD repair was negligible during the first hours following UV irradiation (Extended Data Fig. S1e), indicating that the observed CPD bias was not due to differential repair of UV lesions in daughter cells but rather a bona fide DNA damage segregation bias. Remarkably, this segregation bias was observed both in cancer (U2OS, Fig. 1a) and non-cancer cells (RPE-1, Extended Data Fig. S1f) with a 15% CPD bias on average in both cell types. To investigate whether a DNA segregation bias may explain the asymmetric distribution of UV damage observed in G1 daughter cells, we labeled parental DNA with 5-Chloro-2’-deoxyuridine (CldU)^10^ and measured the parental DNA segregation bias (Extended Data Fig. S1g). As reported before, parental DNA shows a moderate segregation bias between G1 daughter cells (9% on average)^10^ but it did not correlate with the CPD bias observed in the same cell pairs (Extended Data Fig. S1h). This experiment demonstrates that the UV damage distribution bias between daughter cells is not related to a total DNA segregation bias.

**Figure 1:**
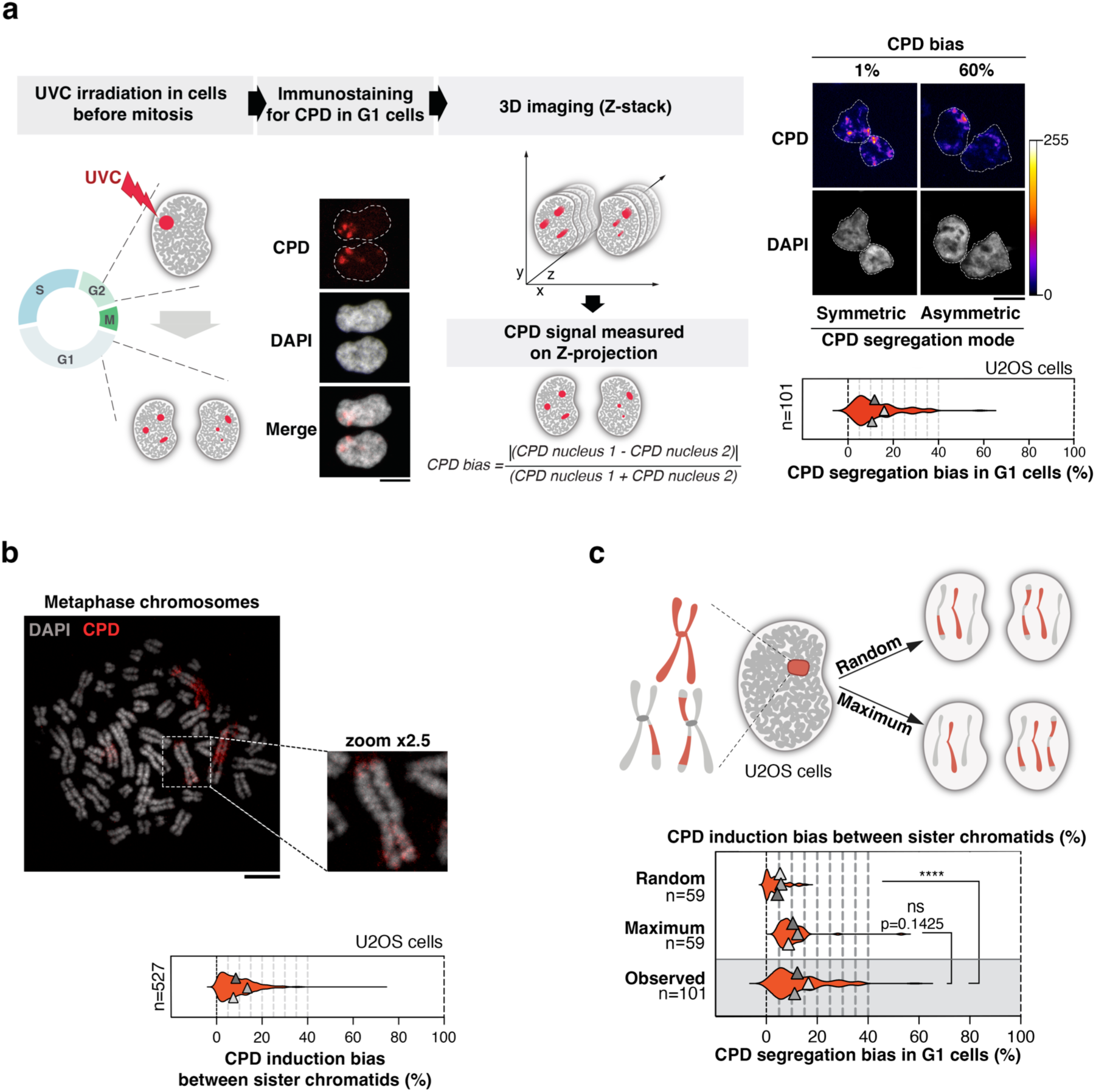
Non-random segregation of UV damage through mitosis. **a.** Left: Experimental procedure for synchronizing cells, inflicting UV irradiation in late G2 and imaging UV damage segregation in G1 cells. Immunofluorescence images show an example of G1 daughter cells harboring scattered UV damage signal (CPD) reflecting the segregation of local UV damage inflicted before mitosis. Right: Pseudo-colored immunofluorescence images (lookup table shown on the right) of CPD distribution between G1 daughter cells (U2OS) to illustrate the symmetric versus asymmetric segregation of local UV damage inflicted before mitosis. The violin plot shows the CPD segregation bias in G1 daughter cells exposed to local UVC irradiation just before mitotic entry. The mean of each independent experiment is represented as a grey triangle ± SD from n cells. The formula to calculate the CPD segregation bias is indicated and further detailed in the method section. Absolute values are used to avoid negative values. **b.** CPD induction bias between sister chromatids analyzed by immunostaining on metaphase spreads from U2OS cells exposed to local UVC irradiation just before mitosis. A representative image is shown with a 2.5x zoom on a damaged chromosome. The CPD bias between each pair of sister chromatids from n damaged chromosomes is represented. **c.** The expected CPD bias in case of random segregation of damaged chromatids (“Random”) and the maximum bias in case of damage-driven chromatid segregation (“Maximum”) are compared to the observed CPD distribution bias in G1 daughter cells shown in panel (a) (“Observed”, grey rectangle). Violin plots show the mean of each independent experiment as a grey triangle ± SD from n cells (a, c) or damaged chromosomes (b). ****: p < 0.0001. Scale bars: 10 µm.

The UV damage distribution bias observed in G1 entails an asymmetrical distribution of UV lesions on sister chromatids. To further dissect this process, we analyzed the distribution of CPDs between sister chromatids in metaphase, which showed close to 10% asymmetry (Fig. 1b). This biased induction of UV lesions between sister chromatids could be due to one sister chromatid partially shielding the other one during the irradiation. Next, we tested if damaged chromatids segregated randomly between daughter cells. Based on the quantification of CPDs on sister chromatids in metaphase nuclei (Fig. 1b), we calculated the CPD bias that would result from a random shuffling of damaged sister chromatids between the two daughter cells (Random) and the maximum segregation bias that we would expect if the most damaged sister chromatids were transmitted to the same daughter cell (Maximum, Fig. 1c). We then compared the results of these simulations to the UV damage bias that we previously observed between G1 daughter cells (Fig. 1a). Interestingly, the observed CPD bias in G1 cells is comparable to the maximum CPD bias and significantly higher than the bias resulting from random DNA damage segregation (Fig. 1c). Together, these results demonstrate that UV-damaged chromatids are segregated through mitosis in a non-random manner.

### Local defect in H3 mitotic phosphorylations at UV damage sites

To dissect the mechanisms that control the non-random segregation of UV-induced damage during mitotic cell division, we first tested whether it was driven by the ATR-Chk1 kinase pathway, as described for the non-random segregation of damaged chromosomes after replication stress ^10^. However, treatment of cells with an ATR inhibitor (ATRi) did not reduce the CPD segregation bias in G1 daughter cells (Extended Data Fig. S1i), suggesting that a different mechanism was involved in response to UV. In line with this, while the DNA damage response at telomeres is, at least partially, involved in the non-random segregation of replication stress-induced damage ^6,10^, we did not observe any significant correlation of the CPD bias with the proportion of telomeric CPD (Extended Data Fig. S1j), arguing that telomeric damage is not driving the asymmetric segregation of UV lesions.

We thus decided to explore other chromatin-based mechanisms that may sustain the non-random segregation of UV-induced damage, focusing on mitotic histone PTMs. One of the most studied histone PTM induced upon mitotic entry is phosphorylation on serine 10 of histone H3 (H3S10ph)^14,15^. We thus monitored the distribution of this mitotic phosphorylation by immunofluorescence in human cells locally irradiated with UV in late G2 and released at the G2/M boundary as above (Extended Data Fig. S1a). We observed a local defect in mitotic H3S10ph at UV sites 1h after UVC irradiation in U2OS, primary BJ and RPE-1 cells (Fig. 2a, Extended Data Fig. S2a-S2c) that correlated with the UV dose (Extended Data Fig. S2d).

**Figure 2:**
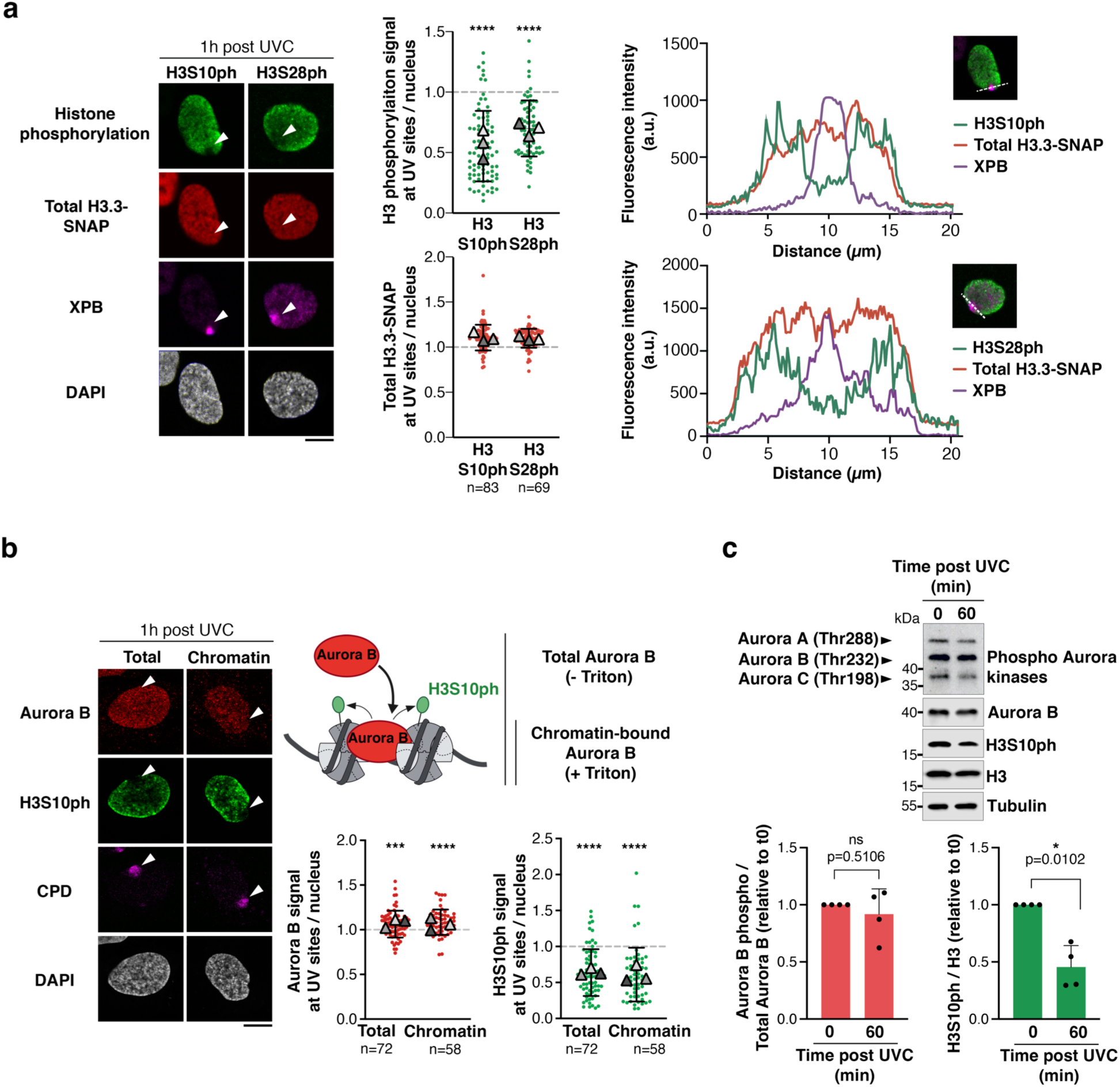
H3 phosphorylation defect at UV damage sites in early mitotic chromatin. **a-b.** H3 phosphorylation (H3S10ph, H3S28ph) and Aurora B kinase localization analyzed at the indicated times after local UVC irradiation in U2OS H3.3-SNAP cells synchronized in early mitosis. The detection of chromatin-bound Aurora B requires detergent permeabilization with Triton-X-100 (+ Triton). Damage sites are marked by XPB or CPD immunodetection and indicated by arrowheads. Dot plots show the mean of each independent experiment as a grey triangle ± SD from n cells (colored dots). The significance of signal enrichment or loss at UV sites is indicated compared to a theoretical value of 1 (dotted line). ****: p < 0.0001. Fluorescence intensity profiles along the dotted lines are displayed on the graphs. **c.** Levels of phosphorylated Aurora B (middle band, Thr232) and H3S10ph monitored by western-blot at the indicated time points post UVC irradiation in U2OS cells (Tubulin, loading control) with corresponding quantifications (mean ± SD from 4 independent experiments). The cell synchronization in G2/M was comparable in all conditions with around 20% of H3S10ph-positive cells scored by immunofluorescence. Scale bars: 10 µm.

Mitotic phosphorylation on serine 28 of histone H3 (H3S28ph) showed a similar defect (Fig. 2a). The reduction in H3S10ph and H3S28ph signals did not result from a local decompaction of chromatin since there was no significant decrease in total H3 density at UV sites, as measured for two different H3 variants (Fig. 2a and Extended Data Fig. S2e). We next examined the kinetics of the H3 phosphorylation defect by irradiating cells at different time points before mitotic entry. These experiments revealed an early and transient defect of H3S10ph in UV-damaged chromatin, restricted to cells that experienced damage within 1.5 h before mitosis (Extended Data Fig. S2f). Despite the decrease in XPB loading on damaged chromatin 3 h after irradiation, the levels of UV damage per se, as assessed by CPD immunodetection, remain stable with time, indicating that the lack of H3S10ph defect at UV sites 3 h after DNA damage induction is not due to the clearance of CPDs (Extended data Fig. S2g). Importantly, the local defect in H3 mitotic phosphorylation at UV damage sites was not explained by the mislocalization or impaired activation of the corresponding kinase Aurora B^16^, whose binding to chromatin (detergent-resistant fraction, Fig. 2b and Extended Data Fig. S2h) and autophosphorylation on threonine 232 (Fig. 2c and Extended Data Fig. S2i) were not significantly inhibited following UV irradiation. These findings uncover a local impairment of mitotic H3 phosphorylations in UV-damaged chromatin that is not due to a defect in the corresponding kinase activity, suggesting that the histone substrate may be unsuitable for the reaction.

### UV-induced chromatin dynamics counteract H3S10ph deposition in UV-damaged chromatin

To characterize histone H3 features impeding its mitotic phosphorylation at UV damage sites, we first explored a potential interference with other H3 PTMs. The detection of neighboring methylation on lysine 9 of histone H3 (H3K9me), after treating fixed cells with lambda-phosphatase to circumvent H3K9 epitope masking by H3S10ph^17^ (Extended Data Fig. S3a), revealed that H3K9me levels remained essentially unaffected at UV sites in early mitotic nuclei, ruling out their contribution to a local inhibition of H3S10ph in response to UV (Extended Data Fig. S3b). Instead, taking advantage of a range of antibodies recognizing H3S10ph in the context of different H3K9me states (Extended Data Fig. S3c)^18^, we noticed that the H3S10ph defect occurred preferentially on H3 devoid of H3K9 tri-methylation (H3K9me3) (detected by the CST and CMA311 antibodies, Extended Data Fig. S3d). This behavior is reminiscent of another H3S10 PTM, the DNA damage-induced ADP-ribosylation (ADPr), which is inhibited by neighboring H3K9me3 and antagonizes H3S10ph^19–21^, hinting towards a possible crosstalk between H3S10 phosphorylation and ADPr by PARP1/2 enzymes (poly(ADP-ribose) polymerases) and their cofactor HPF1 (histone PARylation factor 1) (Fig. 3a). Consistent with this hypothesis and previous reports^22,23^, ADPr was enriched at UV damage sites (Extended Data Fig. S4a). Furthermore, histones represented the major substrates for mono-ADPr following UV irradiation (Extended Data Fig. S4b). Inhibition (Fig. 3b) or knock-down (Extended Data Fig. S4c) of PARP enzymes abrogated the H3S10ph defect in cells damaged just before mitosis, revealing an early inhibitory effect of ADPr on H3S10ph at UV sites. Similarly, knocking-down the PARP cofactor HPF1, that regulates histone ADP-ribosylation in response to DNA damage^24^, reduced H3S10ph defect at UV sites (Fig. S4d), indicating that PARP and HPF1 act in concert to inhibit H3S10ph deposition in UV-damaged chromatin. Reciprocally, maintaining ADPr by inhibiting Poly(ADP-ribose) glycohydrolase (PARG) or by depleting ADP ribosylhydrolase 3 (ARH3), which specifically targets serine residues^19,25^, led to a prolonged H3S10ph defect at UV sites, detectable in cells irradiated 3h before mitosis (Fig. 3c, Extended Data Fig. S4e). To directly and unambiguously address the crosstalk between ADPr and phosphorylation on the H3S10 residue, we introduced in U2OS cells non-ADPribosylatable forms of SNAP-tagged H3.1 or H3.3 by mutating S10 to a threonine (Extended Data Fig. S4f). Importantly, H3T10 can still be phosphorylated in early mitosis, as detected with an antibody that recognizes phosphorylation on both H3S10 and H3T10^26^, albeit with reduced affinity for the threonine mutant (Extended Data Fig. S4g). While a decrease in H3S10ph in response to UV irradiation was observed on endogenous histone H3 and on wild-type SNAP-tagged histones, as expected, the signal was unchanged on non-ADPribosylatable histone mutants (Fig. 3d), suggesting that UV damage-induced ADPr on H3S10 may block the phosphorylation of this residue in early mitotic chromatin. Together, our findings uncover a unique interference mechanism in UV-damaged chromatin where histone ADPr counteracts the deposition of a key mitotic phosphorylation mark on histone H3 at the G2/M transition.

**Figure 3:**
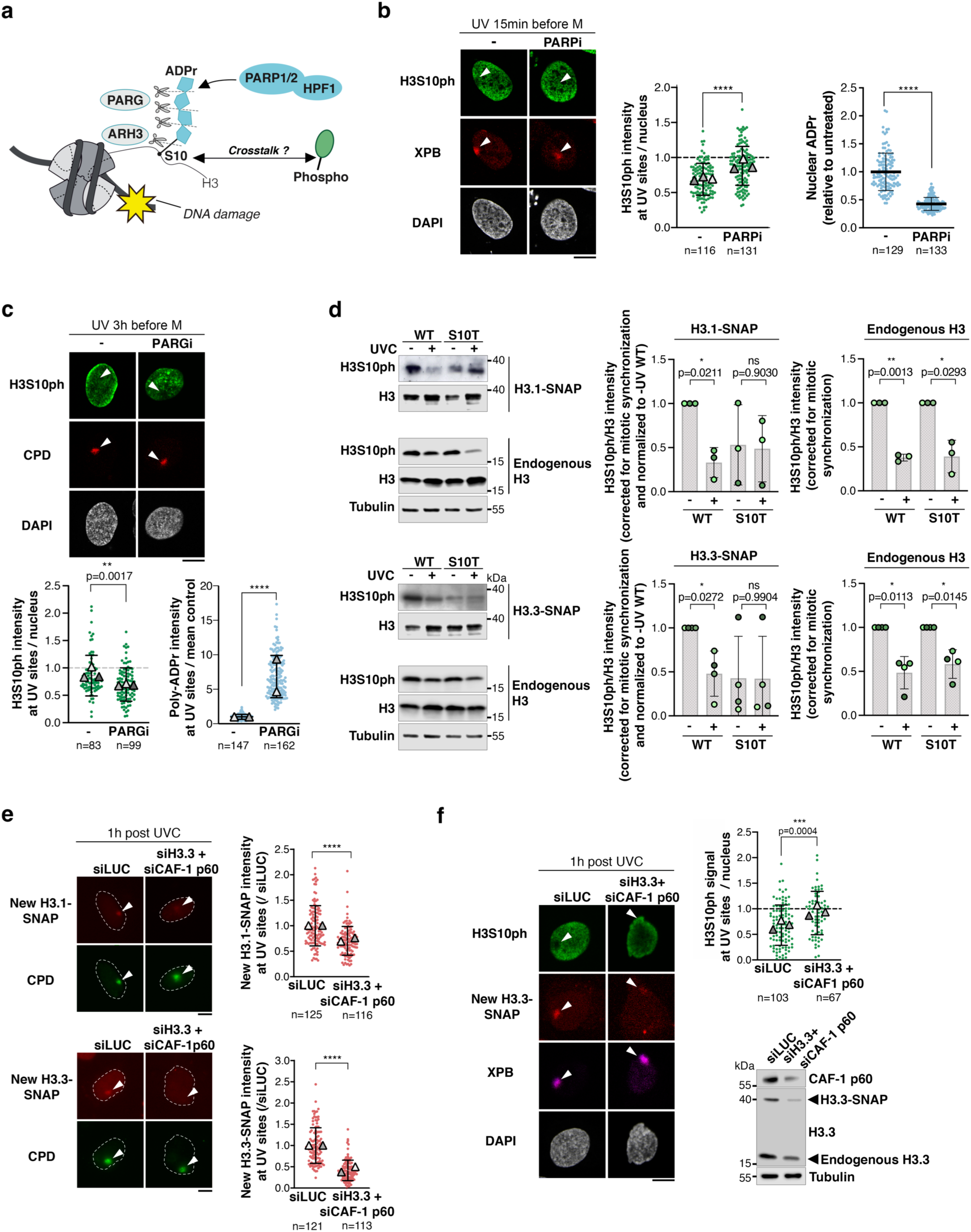
Two independent mechanisms regulate H3S10ph deposition in UV-damaged chromatin. **a.** Working hypothesis for a potential crosstalk between phosphorylation and UV-induced ADP-ribosylation (ADPr) by the PARP-HPF1 axis on H3S10. The contributions of PARG and ARH3 to ADPr removal are indicated. **b-c.** Immunostaining for H3S10ph at UV damage sites (marked by XPB or CPD) in early mitotic U2OS cells treated with the indicated inhibitors (PARPi; PARGi; -, vehicle) and harvested at the indicated times after local UVC irradiation. PARP and PARG inhibitor treatments are controlled by changes in the nuclear poly-ADPr signal measured by immunofluorescence (light blue dot plots). **d.** Levels of H3S10ph monitored by western-blot 30 min post UVC irradiation in U2OS cell lines stably expressing wild-type (WT) or S10T mutated H3.1-SNAP or H3.3-SNAP in a wild-type H3 background (Tubulin, loading control). Quantification of H3S10ph on endogenous and SNAP-tagged H3 is shown on the graphs (mean ± SD from at least 3 independent experiments). All the data were normalized to the endogenous H3S10ph/H3 ratio in non-irradiated conditions in the corresponding cell line to correct for variations in mitotic synchronization, and SNAP-tagged H3 results are presented relative to the -UV WT condition. **e.** New histone H3.1 and H3.3 accumulation at UV damage sites analyzed after local UVC irradiation in asynchronous U2OS H3.1-or H3.3-SNAP cells treated with the indicated siRNAs (siLUC, control). New histones were labeled with SNAP-cell TMR star. **f.** H3S10ph signal analyzed by immunofluorescence after local UVC irradiation in U2OS H3.3-SNAP cells synchronized in early mitosis and treated with the indicated siRNAs (siLUC, control). UV damage sites are marked by XPB immunostaining. Knock-down efficiencies are controlled by western-blot (Tubulin, loading control). Dot plots show the mean of each independent experiment as a grey triangle ± SD from n cells (colored dots). Arrowheads point to UV-damaged regions. ****: p < 0.0001. Scale bars: 10 µm.

### New H3 deposition at UV sites delays mitotic H3S10ph

As the effect of histone ADPr on H3S10ph was restricted to cells damaged just before mitosis (observed 15 min before, Fig. 3b, and not 1h before, Extended Data Fig. S5a), we searched for additional mechanisms that could contribute to the H3S10ph defect after the hydrolysis of ADPr chains. Newly synthesized H3 is incorporated at sites of UV damage repair^27,28^ and displays less phosphorylation on S10 in early mitosis compared to old histones^29^. Newly synthesized H3 may thus be a poor substrate for mitotic phosphorylation, delaying H3S10ph deposition at UV sites. Interfering with new H3 deposition pathways by treating cells with siRNAs against H3.3 and the H3.1-specific chaperone CAF-1 (Chromatin Assembly Factor-1) blocked new H3 incorporation at UV sites (Fig. 3e) and abolished the local defect in H3S10ph at UV sites in early mitotic cells (Fig. 3f). Importantly, blocking new H3 deposition did not generate UV damage on its own, which could have impaired H3S10ph levels (Extended Data Fig. S5b). In contrast to the dual depletion of H3.3 and CAF-1, single depletions of H3.3, of the H3.3 chaperone HIRA (Histone Regulator A), which is the main chaperone involved in new H3.3 deposition at damage sites^30,31^, or of the H3.1 chaperone CAF-1, had only a moderate effect on H3S10ph at UV sites (Extended Data Fig. S5c-e). These results indicate that *de novo* deposition of both H3 variants redundantly controls mitotic H3S10ph in UV-damaged chromatin. As an orthogonal approach, we interfered with new H3.1 and H3.3 deposition at UV sites by depleting the DNA damage sensor DDB2 (DNA damage binding protein 2) (Extended Data Fig. S5f), which also restored H3S10ph at UV damage sites (Extended Data Fig. S5g). This pathway appears to be distinct from the PARP-mediated mechanism described above since ADPr at UV sites was independent of DDB2 (Extended Data Fig. S5h). Thus, our findings reveal two independent layers of control, histone ADPr and new H3 deposition, both interfering with mitotic H3S10ph in UV-damaged chromatin.

### The H3S10ph control pathway governs UV damage segregation in daughter cells

To test whether mitotic H3S10ph controls the DNA damage segregation process, we measured CPD levels in G1 daughter cells in two conditions showing opposite effects on H3S10ph at UV sites in early mitosis: UV irradiation 3h before release into mitosis, which does not lead to a detectable H3S10ph defect, vs. UV irradiation immediately before release into mitosis leading to an H3S10ph defect (Extended Data Fig. S2f). Remarkably, cells irradiated just before release into mitosis displayed a stronger bias of UV damage segregation between daughter cells in U2OS (Fig. 4a). We obtained similar results in RPE-1 cells (Extended Data Fig. S6a) indicating that this phenomenon was not restricted to cancer cells. As the CPD signal remained unchanged throughout the experiment (Extended Data Fig. S1e), differences in segregation bias were not due to variable amounts of UV damage. The observed differences in segregation bias also did not relate to differences in damage distribution between sister chromatids, which were comparable between cells irradiated 3h and just before release into mitosis (Extended Data Fig. S6b). Mechanistically, restoring H3S10ph by PARP inhibition significantly reduced the CPD segregation bias in daughter cells (Fig. 4b and Extended Data Fig. S6c). Reciprocally, PARG inhibition, which prolongs the H3S10ph defect in early mitosis, resulted in an increased CPD segregation bias in G1 daughter cells (Fig. 4b and Extended Data Fig. S6d). Note that PARP inhibition was applied to cells in late G2, thus avoiding the adverse mitotic consequences of PARP inhibition in S phase^32^. We attribute the result of PARP inhibition to the H3S10ph defect rather than to a distinct PARP-dependent process since the PARPi treatment on cells damaged 1h before release into mitosis did not interfere with H3S10ph deposition at UV sites (Extended Data Fig. S5a) and had no effect on DNA damage distribution between G1 cells (Extended Data Fig. S6e). Strengthening these findings, depletion of the DNA damage sensor DDB2, which also abrogates the H3S10ph defect, similarly reduced the UV damage segregation bias (Fig. 4c). We conclude that the H3S10ph control pathway that marks damaged chromatin in early mitosis governs the asymmetric segregation of UV damage in daughter cells. UV damaged chromosomes fail to properly condensate in early mitosis as shown by a lower coefficient of variation of DAPI staining at sites of local UV irradiation (Extended data Fig. S6f), which might interfere with the segregation of the damaged genetic material. However, the chromosome condensation defect at UV sites was unaffected by PARP inhibition (Extended Data Fig. S6g). This indicates that the lower condensation of UV-damaged chromosomes is unlikely to contribute to the asymmetric segregation process, which is PARP-dependent.

**Figure 4:**
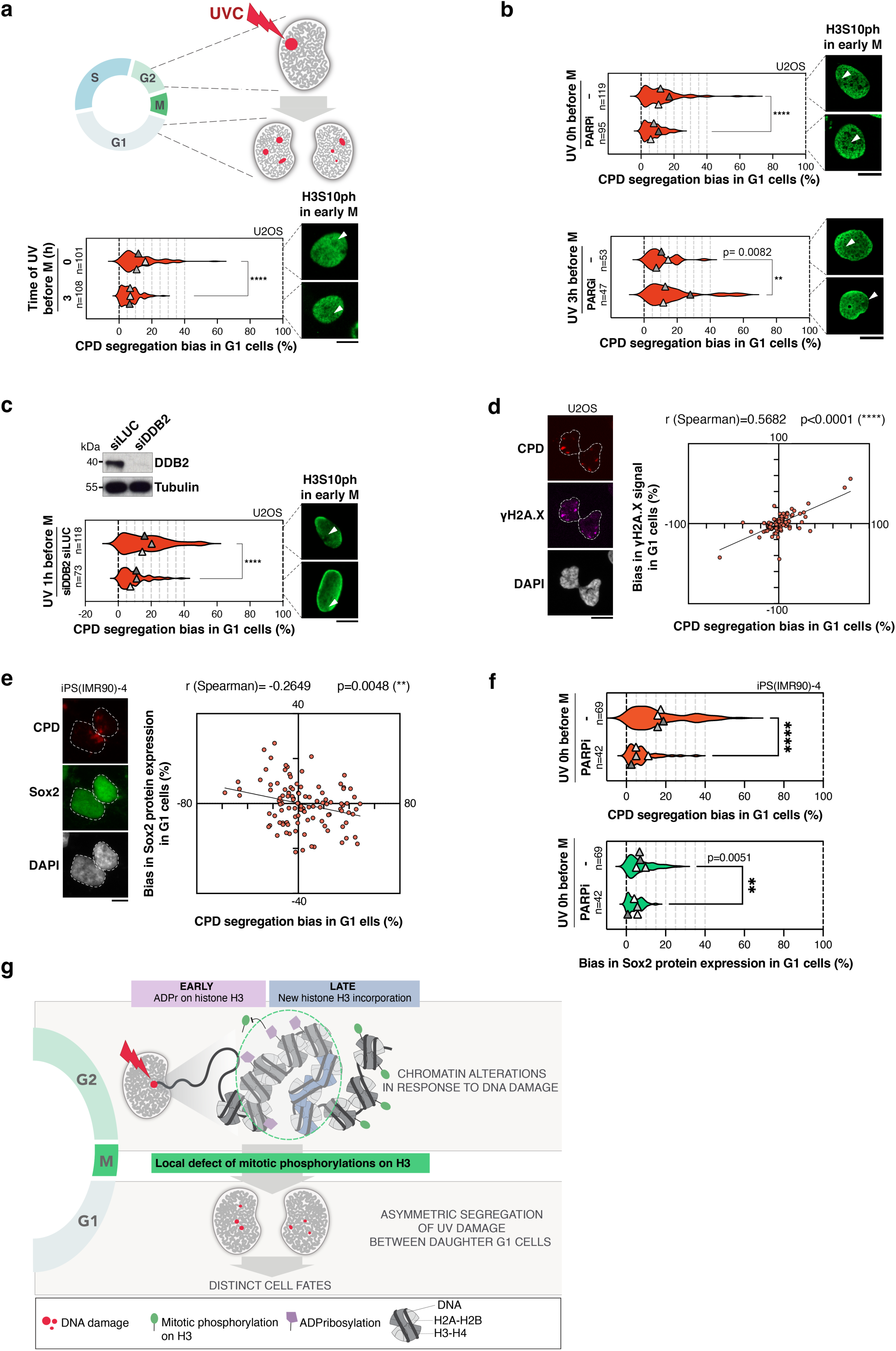
The UV-dependent H3S10ph control pathway biases DNA damage segregation in mitosis. **a-c**. Segregation of the DNA damage marker CPD assessed by immunostaining in G1 daughter cells (U2OS) treated with PARP or PARG inhibitors (**b**; -, vehicle) or depleted for the DNA damage sensor DDB2 (**c**; siLUC, control) and irradiated at the indicated time points before mitosis. Immunofluorescence images illustrate the corresponding H3S10ph pattern at UV sites (white arrowheads) in early mitosis for each condition. Knock-down efficiency is controlled by western-blot (Tubulin, loading control). **d.** Correlation analysis between the UV damage segregation bias (CPD) and the bias in γH2A.X signal in U2OS G1 daughter cells (102 cell pairs were scored in 3 independent experiments). **e.** Correlation analysis between the UV damage segregation bias (CPD) and the bias in Sox2 protein expression in iPS(IMR90)-4 G1 daughter cells (112 cell pairs scored in 5 independent experiments). **f.** Effect of PARP inhibition (-, vehicle) on the UV damage segregation bias (CPD) and on the bias in Sox2 protein expression in iPS(IMR90)-4 G1 daughter cells. Violin plots show the mean of each independent experiment as a grey triangle ± SD from n cells in total. **g.** Working model depicting the two layers of control that antagonize H3S10ph deposition at UV damage sites in early mitosis and their consequences on DNA damage segregation in daughter cells. ****: p < 0.0001. Scale bars: 10 µm.

We further analyzed the consequence of the asymmetric distribution of UV damage on downstream DNA damage signaling, which involves phosphorylation of the histone variant H2A.X^33^ . We observed a bias in γH2A.X signal between G1 daughter cells, which significantly correlated with the CPD segregation bias (Fig. 4d). This suggests that the asymmetric segregation of UV damage governs the strength of DNA damage signaling in daughter cells. Propagation of DNA damage to daughter cells would be particularly deleterious for stem cell integrity^34^, which prompted us to investigate the consequence of asymmetric DNA damage segregation on stemness properties. We took advantage of the ability of human induced pluripotent stem cells (iPSC) derived from IMR90 fibroblasts to generate daughter cells with different fates, one retaining stem cell properties while the other commits to differentiation. Of note, iPSCs are equipped to repair a large variety of DNA damage including UV lesions^35^. We examined the expression of the stem cell marker Sox2 to evaluate the impact of UV lesion segregation on the stemness status post mitosis. We noticed a moderate bias in Sox2 protein expression levels between daughter cells, which showed a weak but significant anticorrelation with the CPD segregation bias (Fig. 4e). This result indicates that the daughter cells inheriting the most damaged chromosomes tend to have reduced stemness capacity. Furthermore, PARP inhibition diminished both the CPD segregation bias as observed in U2OS cells (Fig. 4b) and the bias in Sox2 expression levels between damaged daughter cells (Fig. 4f). Notably, despite the functional interaction between PARP1 activity and Sox2 protein levels^36^ short-term PARP inhibition (2.5 h in total in our experiments) had no significant impact on overall Sox2 levels (Extended Data Fig. S6h). These results suggest that stem cells may be protected from the transmission of high levels of DNA damage through the asymmetric segregation of DNA lesions during mitotic cell divisions.

## DISCUSSION

Understanding how chromatin integrity and cell identity are preserved from deleterious genotoxic stress through cell divisions is of fundamental importance. By studying mitotic phosphorylations on histone H3 in response to UV damage, our work sheds light on a chromatin-based mechanism that governs the non-random segregation of DNA lesions (Fig. 4g). The discovery that chromatin alterations may impinge on genome stability not only by regulating DNA repair but also by controlling DNA damage segregation in mitosis broadens the scope of genome maintenance mechanisms.

We identify two complementary pathways that interfere with mitotic H3S10ph in UV-damaged chromatin, with an early contribution of histone ADP-ribosylation and a later contribution of newly deposited histone H3 (Fig. 4d). Our findings support a model, distinct from what was recently shown for replication stress-induced damage^6,10^, where the resulting H3S10ph defect may drive the asymmetric segregation of UV-damaged chromatin in daughter cells, with potential consequences on cell fate.

Our study also provides important insights into the so far elusive roles of histone ADPr and new histone deposition at DNA damage sites. We demonstrate that they both contribute to bookmark damaged chromatin during a crucial stage of the cell cycle by interfering with H3 mitotic phosphorylation. This function of histone ADP-ribosylation is likely distinct from its recently reported role in promoting chromatin relaxation at DNA double-strand breaks in human cells^24^. We provide evidence for a direct interference between ADPr and phosphorylation on H3S10 and the same likely applies to H3S28 because those two residues are the main targets of damage-induced ADPr^21^. Notably, a similar interplay between H3S10/28 ADPr and phosphorylation was recently reported in response to DNA double-strand breaks in amoeba, impacting mitotic entry of cells with unresolved damage^26^. Regarding newly synthesized H3, we suspect that they impair H3S10ph because they may represent poor substrates for Aurora B due to their specific PTM pattern. Indeed, Aurora B preferentially phosphorylates S10 on hypoacetylated H3 tails^37^, while new H3 is enriched for acetylation on lysines 14 and 23^38^. We note that besides these effects at the level of the histone substrate, we cannot rule out a possible contribution of phosphatases in H3S10ph removal from UV-damaged chromatin, as reported in G1 cells after ionizing radiation-induced damage^39^. We did not notice any negative impact of UV damage on the Aurora B kinase in contrast to the PARP-dependent inhibition of Aurora B reported following oxidative damage^40^.

A major conclusion of our work is the characterization of non-random segregation of post-replicative UV damage in the cell progeny. We propose that the local alteration in mitotic H3 phosphorylation could drive the asymmetric segregation of DNA damage, thus impacting daughter cell fate. An analogy can be drawn with another mitotic phosphorylation on H3T3, which governs asymmetric segregation of histone proteins and daughter cell fates in the Drosophila male germline^13^. Similar to H3S10ph, H3T3ph distinguishes pre-existing from newly synthesized histones. H3T3ph drives the asymmetric segregation of old and new histones in daughter cells because old and new histones H3 are asymmetrically distributed on the two strands during replication in Drosophila^41,42^, while in mammalian cells they distribute symmetrically^43^. We propose that the induction of post-replicative UV damage in human cells may generate a local asymmetry at the histone level between sister chromatids, allowing the asymmetric segregation of DNA damage in daughter cells. How the H3S10ph defect at UV sites is mechanistically connected to the asymmetric distribution of damaged chromosomes is a pending question. This may rely on an asymmetry in the chromosome segregation machinery with asymmetric microtubule assembly and attachment to kinetochores, as described in Drosophila^44,45^. A local lack of histone phosphorylation may also affect the electrostatic properties of mitotic chromatin, known to control chromosome separation^46,47^, which would favor the co-segregation of damaged chromatids. Furthermore, the chromatin context where UV damage occurs may modulate DNA damage inheritance through cell divisions.

We speculate that the damaged material would be preferentially inherited by differentiated cells, shown to repair UV damage more efficiently^48^, thus preserving the stem cell compartment. This is particularly important in stem cells, which are devoid of classical cell cycle checkpoints, and would contribute to control genome stability at the tissue scale. By studying asymmetric divisions of human iPSC, we demonstrated that the daughter cells inheriting more UV damage tend to lose stemness potential, and may preferentially adopt a differentiated fate, consistent with the fact that the DNA damage response promotes cell differentiation^49^. Longer term kinetic studies will be needed to further investigate the switch in expression between stemness and differentiation markers upon asymmetric DNA damage segregation in stem cell populations. Given that we also observed a UV damage segregation bias in U2OS and RPE-1 cells, which do not switch between stem cell and differentiated states upon division, the non-random segregation of UV damage is more likely a cause than a consequence of distinct daughter cell fates. In addition, daughter cells inheriting more UV damage also exhibit stronger DNA damage signaling, highlighting multiple complementary strategies to preserve genome integrity post mitosis. An interesting possibility that deserves further investigation is that the non-random segregation of UV damage may be accompanied by an asymmetric distribution of repair proteins, reflecting distinct repair capacities in daughter cells, as described following replication stress^6,10^. Studying the damage response in physiological models that harbor asymmetric cell divisions will be of high interest to further investigate the functional consequences of non-random damage segregation on cell identity and stem cell homeostasis.

## METHODS

### Cell culture and drug treatments

BJ primary fibroblasts (ATCC CRL-2522, human foreskin, male), U2OS cells (ATCC HTB-96, human osteosarcoma, female) and RPE-1 hTERT cells (ATCC CRL-4000, human retinal pigment epithelium, female) were grown at 37°C and 5% CO_2_ in Dulbecco’s modified Eagle’s medium (DMEM, Invitrogen) supplemented with fetal calf serum (EUROBIO, 15% final for BJ and 10% final for U2OS and RPE-1) and antibiotics (100 U/ml penicillin, 100 µg/ml streptomycin, Invitrogen). U2OS cells stably expressing H3.1-SNAP, H3.3-SNAP, and GFP-XPC were grown in the same medium supplemented with selection antibiotics (Supplementary Table 1). IMR90-derived human induced pluripotent stem cells iPS(IMR90)-4 (WiCell) were cultured on Matrigel-coated substrate (Corning® Matrigel® Basement Membrane Matrix) in mTeSR Plus medium (STEMCELL Technologies) supplemented with 0.1% Normocin (Invivogen) at 37°C and 5% CO_2_. The medium was supplemented with 10 µM Y-27632 (ROCK inhibitor, Miltenyi) for the thawing and freezing processes only. The ATR inhibitor AZ20 (Euromedex), and the PARP inhibitor KU-58948 (Axon MedChem) were added to the culture medium at 2 µM and 10 µM final concentration, respectively, 1 h prior UV irradiation. Cells were treated overnight with the PARG inhibitor (Sigma-Aldrich) at 1 µM final concentration. Oxidative damage was induced by treating cells for 10 min with 2 mM hydrogen peroxide (H_2_O_2_, Sigma-Aldrich).

**Supplementary Table 1.** Stable cell lines. Antibiotics: Hygromycin, G418 (Euromedex).

| Stable cell lines | Selection antibiotics | References |
| --- | --- | --- |
| U2OS H3.1-SNAP | 100 µg/ml G418 | 50 |
| U2OS H3.3-SNAP | 100 µg/ml G418 | 50 |
| U2OS H3.3-SNAP GFP-XPC | 100 µg/ml G418 + 200 µg/ml Hygromycin | 51 |

### Site directed mutagenesis

The mutation of Serine 10 to Threonine (S10T) was introduced by site directed mutagenesis on C-terminal SNAP-tagged H3.1 (HIST1H3C coding sequence cloned in pSNAPm) and H3.3 (H3F3B coding sequence cloned in pSNAPm) encoding plasmids with a PCR master kit (Roche) and the primers indicated in Supplementary table 2. U2OS cell lines stably expressing the mutated H3.1 or H3.3 S10T were generated by transfection of the plasmids with Lipofectamine 2000 (Invitrogen). The clones were selected in medium supplemented with G418 (Life Technologies) starting 48h after transfection. Cells were grown in the same conditions as described above for U2OS cells expressing wild-type H3.1-SNAP or H3.3-SNAP. Clones were selected based on total levels of SNAP-tagged histones (see section SNAP labeling of histone proteins) compared to U2OS cells expressing wild-type H3.1-SNAP or H3.3-SNAP.

**Supplementary Table 2.**
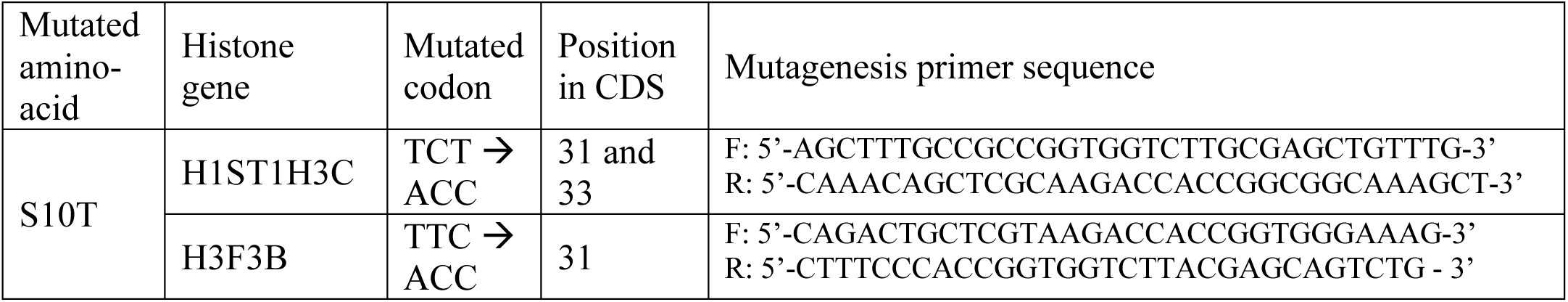
Primers for site directed mutagenesis. CDS: coding sequence; F: forward; R: reverse.

### Cell synchronization

Cell cultures were enriched in early mitotic cells by overnight treatment with 2 mM Thymidine (Sigma-Aldrich), followed by a 6-8h release in fresh medium, and overnight treatment with 10 µM of the Cdk1 inhibitor RO-3306 (Merck Millipore). Cells arrested in late G2 were then allowed to enter mitosis with a 15-min release in fresh medium. To harvest cells in G1 instead of early mitosis, cells were released from the Cdk1 inhibitor for 1h30 in fresh medium.

### UVC irradiation

Local UVC irradiation was performed on cells grown on glass coverslips (VWR) covered with a polycarbonate filter (5 µm pore size, Merck Millipore) and irradiated with UVC (254 nm) using a low-pressure mercury lamp (Vilber-Lourmat). The UVC dose (300 J/m^2^ unless stated otherwise) was controlled using a VLX-3W dosimeter (Vilber-Lourmat). iPS(IMR90)-4 were grown on matrigel (Corning® Matrigel® Basement Membrane Matrix)-coated glass coverslips and irradiated with UVC as described above. The average area of UV damage spots is 23 μm^2^, based on CPD immunostaining, which is consistent with the filter pore size (expected damage area 20 μm^2^). For global irradiation, cells in PBS (Phosphate Buffer Saline) were exposed to 10 J/m^2^ UVC.

### SNAP labeling of histone proteins

Total SNAP-tagged histones were labeled by incubating cells with 2 µM of the red-fluorescent reagent SNAP-cell TMR-star (New England Biolabs) for 15 min before cell fixation. For specific labeling of neo-synthesized histones, pre-existing SNAP-tagged histones were quenched during 30 min with 10 µM of the non-fluorescent reagent SNAP-cell Block (New England Biolabs), followed by a 30-min wash in fresh medium and a 2-h chase. New histones synthesized during the chase step were labeled by incubating cells with the red-fluorescent SNAP reagent as described above or with SNAP-cell Oregon green at 4 µM (15-min pulse), followed by 30 min incubation in fresh medium. Local UVC damage was inflicted just before the pulse.

### Labeling of total parental DNA with CldU

U2OS cells were grown for 2 weeks in culture medium containing 1 μM 5-Chloro-2’-deoxyuridine (CldU, Sigma-aldrich) with medium renewal every 72 h to label all parental DNA and then cultured for 2 days (2 cell divisions) in CldU-free medium in parallel with cell synchronization in G2/M. After local UVC irradiation and release through mitosis into G1, cells were fixed with 2% paraformaldehyde for 15 minutes followed by permeabilization in PBS containing 0.5% Triton-X-100 and denaturation with HCl 2N containing 0.5% Triton-X-100 for 1 h at room temperature. CldU was detected with an anti-BrdU antibody (Supplementary Table 4) following the immunofluorescence procedure.

### siRNA transfection

siRNA purchased from Eurofins MWG Operon or Sigma-Aldrich (Supplementary Table 3) were transiently transfected into cells using Lipofectamine RNAiMax (Invitrogen) at a final concentration of 25-60 nM in the culture medium following manufacturer’s instructions. Cells were harvested 48-72 h after siRNA transfections or 24 h for siH3.3 alone.

**Supplementary Table 3.**
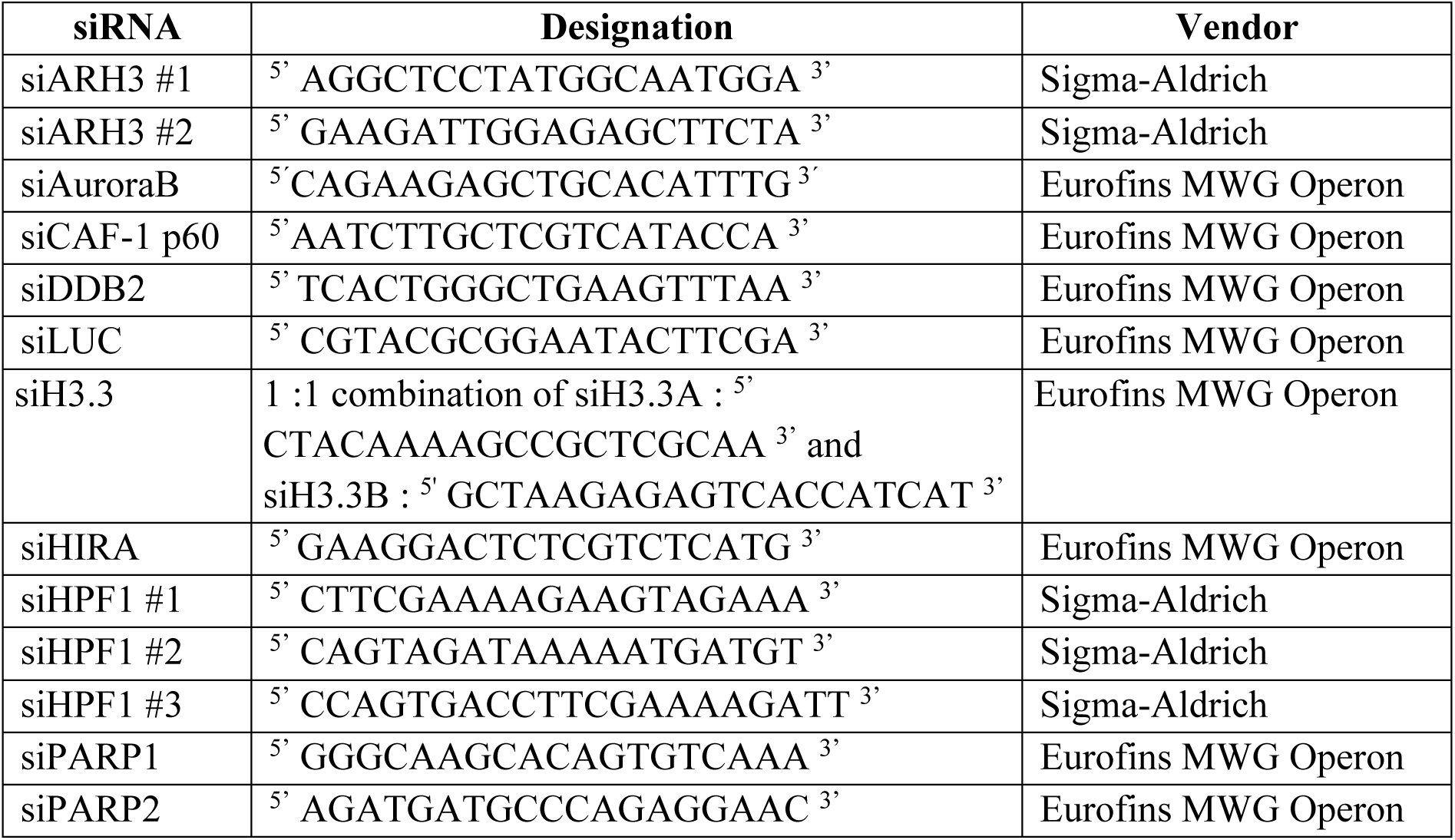
siRNA sequences.

### Immunofluorescence

Cells grown on glass coverslips (VWR) were fixed with 2% paraformaldehyde for 15 min and then permeabilized in 0.2% Triton-X-100 in PBS. For SNAP labeling experiments on asynchronous cell populations and for the detection of chromatin-bound Aurora B, cells were subject to detergent-permeabilization with 0.5% or 0.2% Triton-X-100, respectively, in CSK buffer (Cytoskeletal buffer: 10 mM PIPES pH 7.0, 100 mM NaCl, 300 mM Sucrose, 3 mM MgCl_2_) for 5 min before fixation (pre-extraction). When performed on G2/M synchronized samples, pre-extraction conditions were milder with 0.1% Triton-X-100 in CSK buffer for 1 min to preserve early mitotic cells. For CPD staining, samples were denatured with 0.5 M NaOH for 5 min after fixation. To reveal H3K9 epitopes in early mitotic nuclei, cells were treated after fixation with λ-phosphatase (400 units final, New England Biolabs) for 30 min at 37°C as indicated in the manufacturer’s instructions^17^. After fixation and permeabilization, samples were blocked in 5% BSA (Bovine Serum Albumin, Sigma-Aldrich) in PBS supplemented with 0.1% Tween 20 (Euromedex) and then incubated sequentially with primary antibodies and secondary antibodies conjugated with Alexa Fluor dyes and diluted in blocking buffer (Supplementary Table 4). The choice of antibodies to detect UV damage sites, either against CPD or against the DNA repair factor XPB, is based on the timing post irradiation, the CPD signal being more stable over time, and on the species of other antibodies used in combination. For CPD and Sox2 co-detection, CPD detection with primary and secondary antibodies was performed first as described above, followed by an overnight incubation at 4°C with DyLight 488-coupled Sox2 antibody. Coverslips were mounted in Vectashield or Vectashield plus antifade medium with DAPI (Vector Laboratories).

**Supplementary Table 4:**
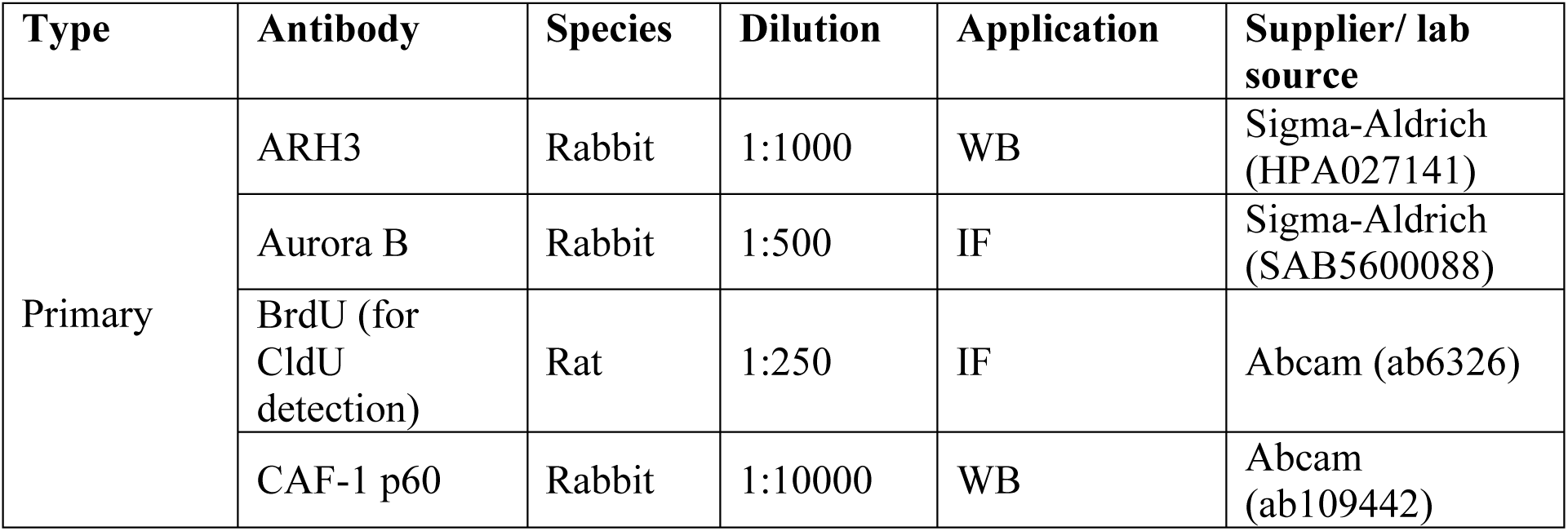
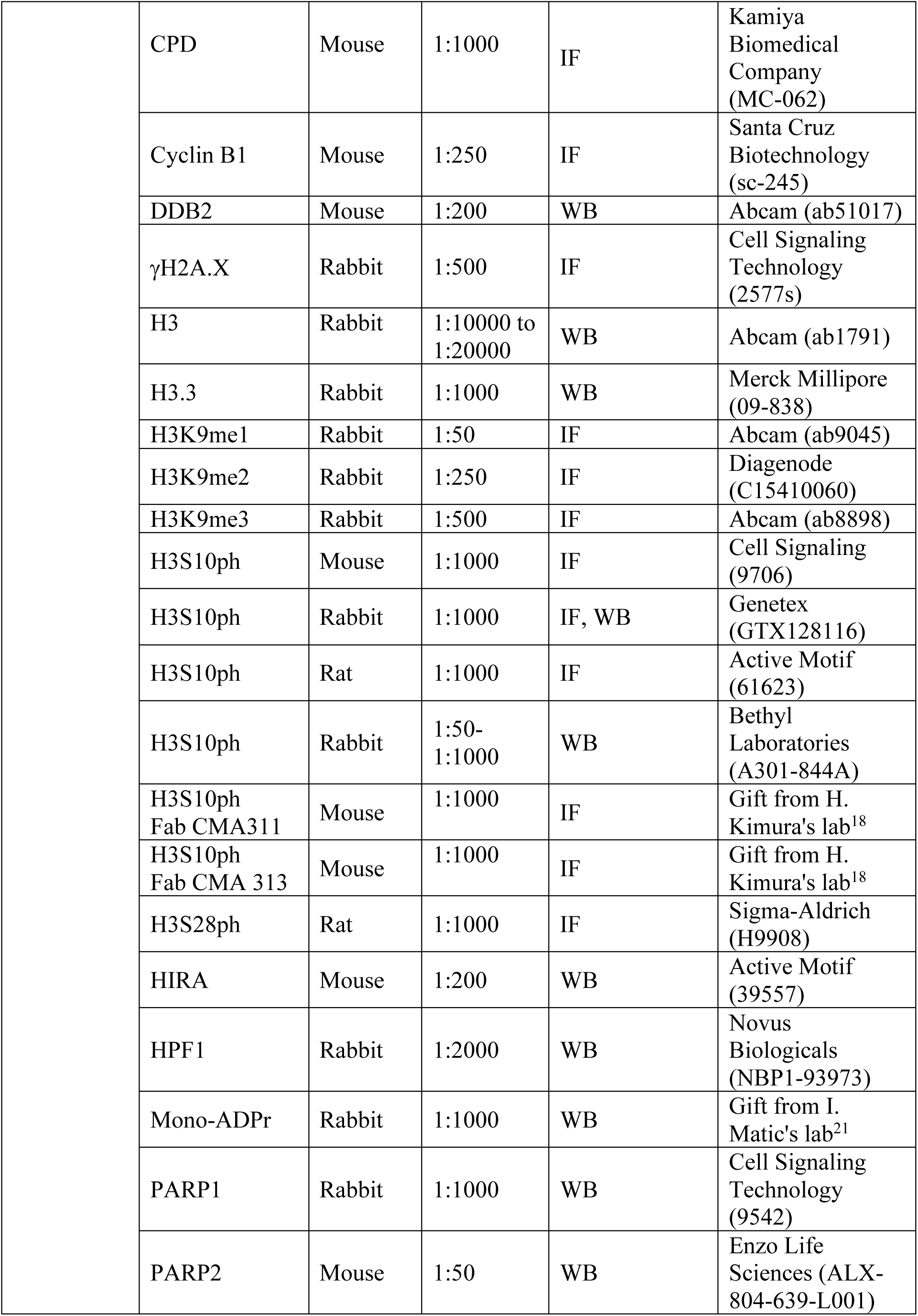
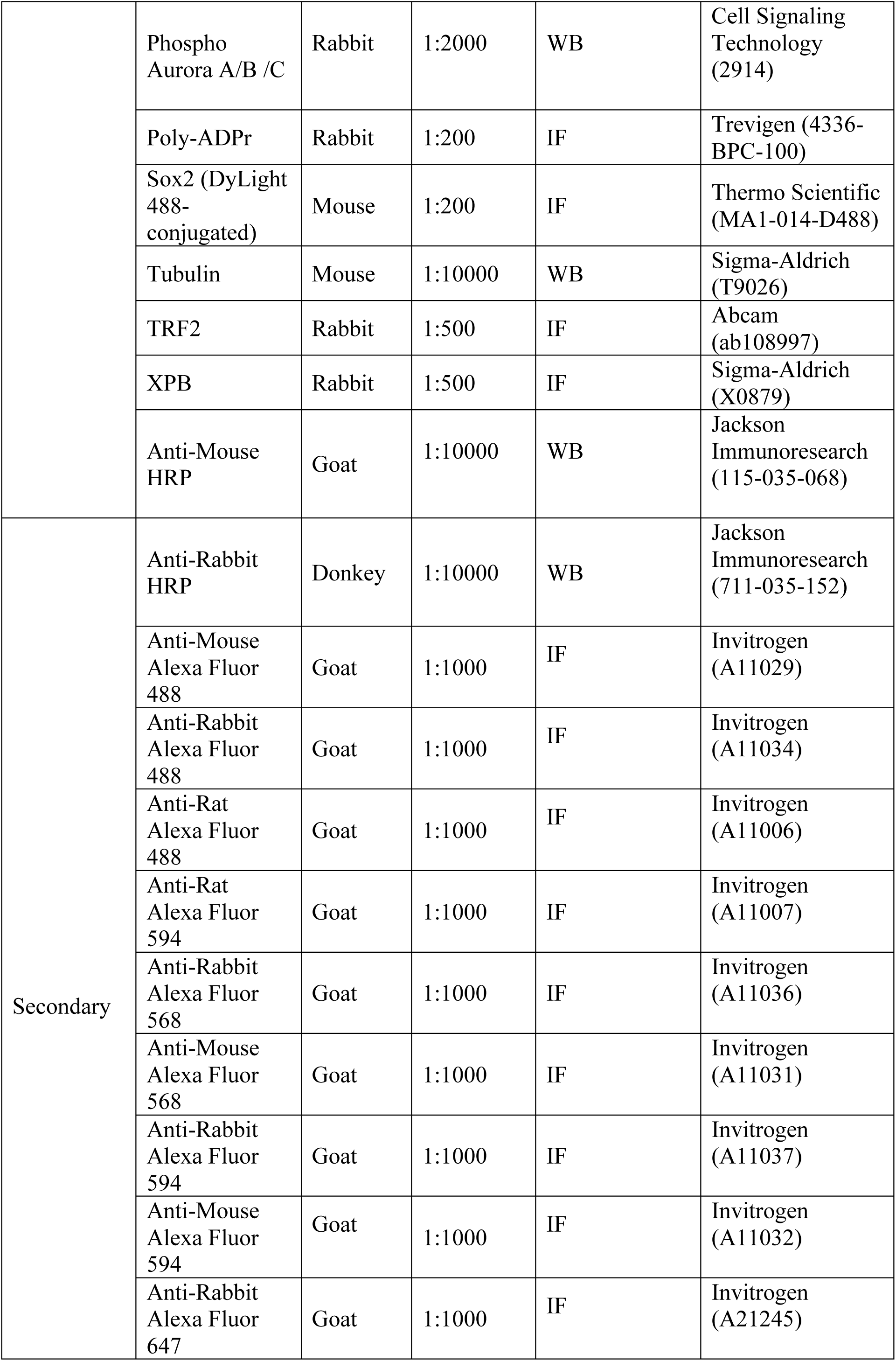
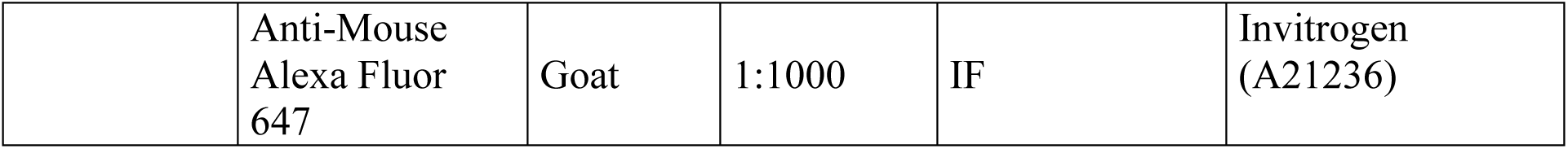
Antibodies. IF: Immunofluorescence; WB: Western-Blot

### Specificity of H3S10ph antibodies

The specificities of H3S10ph antibodies used in this study (Supplementary Table 4) were tested on a histone peptide microarray as previously described^52^. Briefly, chemically synthesized C-terminally biotinylated histone H3 peptides bearing the modifications of interest were spotted onto streptavidin-coated slides using an Aushon 2470 microarrayer. Each peptide was printed as 2 rows of 3 consecutive spots. Slides were hybridized with primary antibodies (CST 9706 1:1000, GTX 128116 1:2000, Abcam ab14955 1:1000, Bethyl A301-844A 1:1000) and Alexa Fluor 647-labeled secondary antibodies (Invitrogen A-21245, A-21235, A-21247, all at 1:5000 dilution). Hybridization steps were performed in buffer containing 25 mM HEPES pH 7.5, 100 mM NaCl, 0.5% BSA (w/v) and 0.1% NP-40 and washes in PBS supplemented with 0.1% Tween 20. Fluorescent signals were read on Innopsys InnoScan 1100AL microarray scanner and analyzed using ArrayNinja^53^. Values were normalized to the maximum signal on the array, and results are presented as mean values +/-SD from 6 spots/peptide.

### Metaphase spreads

U2OS were synchronized in late G2 and damaged locally with the UVC lamp as described above prior to release in metaphase in medium containing 100 ng/ml Colcemid (Sigma-Aldrich) for 2h at 37°C. Cells were trypsinized, resuspended in a 75 mM KCl hypotonic solution for 15 min at 37°C and fixed with ethanol/acetic acid (v:v = 3:1) overnight at -20°C. The next day, the cells were dropped onto glass slides and air dried overnight. The cells on slides were fixed with 2% PFA and processed for immunostaining against the UV damage marker CPD (see Supplementary Table 4) as described above.

### Cell extracts and western-blot

Total extracts were obtained by scraping cells in Laemmli buffer (50 mM Tris HCl pH 6.8, 1.6% Sodium Dodecyl Sulfate, 8% glycerol, 4% β-mercaptoethanol, 0.0025% bromophenol blue) followed by 5 min denaturation at 95°C. Total protein concentration was measured on a DS-11 FX+ spectrophotometer (DenoVix).

Extracts were run on 4-20% MINI-PROTEAN TGX gels (Bio-Rad) in running buffer (200 mM glycine, 25 mM Tris, 0.1% SDS) and transferred onto nitrocellulose membranes (Protran) with a Trans-blot SD semidry transfer cell (Bio-Rad). Total proteins were revealed with Pierce reversible stain (Thermo Scientific). Proteins were probed with the appropriate primary antibodies and Horse Radish Peroxydase (HRP)-conjugated secondary antibodies (Supplementary Table 4). Protein detection by chemiluminescence was achieved using SuperSignal West Pico or Femto chemiluminescence substrates (Pierce) on hyperfilms MP (Amersham) processed in a SRX-101A tabletop film processor (Konica Minolta) or using the Odyssey Fc-imager (LI-COR Biosciences) following manufacturer’s instructions. Quantification of protein levels was performed on images from the Odyssey Fc-imager with the Image Studio Lite software (version 5.2.5).

### Flow cytometry

Cells were fixed in ice-cold 70% ethanol before DNA staining with 50 µg/ml Propidium Iodide (Sigma-Aldrich) in PBS containing 0.05% Tween 20 and 0.5 mg/ml RNaseA (USB/Affymetrix). DNA content was analyzed by flow cytometry (20,000 cells per condition) on a FACS Calibur Flow Cytometer (BD Biosciences) and data were processed with the FlowJo 9.7.5 software (Treestar) using both Watson and Dean/Jett/Fox algorithms for cell cycle analysis.

### Image acquisition and analysis

Confocal images were captured on Zeiss LSM710/780/900 microscopes equipped with a Plan Apochromat 63x/1.4 and 40x/1.3 oil objective and Zen software. Epifluorescence imaging was performed on a Leica DMI6000 microscope using a Plan Apochromat 40x/1.3 or 63x/1.4 oil objective. Images were captured using a CCD camera (Photometrics) and Metamorph software. Images were assembled with Adobe Photoshop. Unprocessed images were analyzed with Image J software (U. S. National Institutes of Health, Bethesda, Maryland, USA, http://imagej.nih.gov/ij/, version *v.2.0.0-rc-69/1.52.p*) after removal of background fluorescence signal. Custom-made Image J macros were used for fluorescence quantifications in the damaged regions (defined by CPD signal or repair factor staining with the “threshold” and “convert to mask” functions) and in the entire nucleus (delineated based on DAPI staining). The enrichment/loss of proteins or histone marks at UV damage sites was given by the ratio of their mean intensities at UV sites relative to the whole nucleus.

The UV damage segregation bias between daughter cells was calculated as the absolute difference of CPD intensities divided by the sum of CPD intensities measured in daughter cells as indicated below (cell1 and cell2 were defined arbitrarily among the pair of G1 cells). The resulting value was then converted into percent.

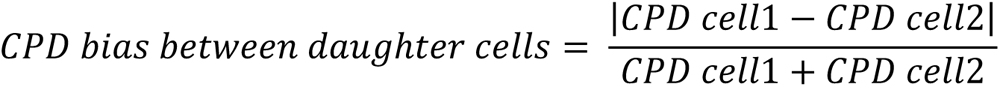

When correlating the CPD bias with another parameter measured in daughter cells (e.g. CldU, γH2A.X or Sox2 signal), the same formula as above was used to calculate intensity biases between daughter cells, except using the relative difference instead of the absolute difference. For CPD bias calculations on metaphase spreads, the CPD intensity was measured on each sister chromatid in all UV-damaged chromosomes and the CPD bias was calculated as follows (sister1 and sister2 were defined arbitrarily among the pair of sister chromatids) and converted into percent:

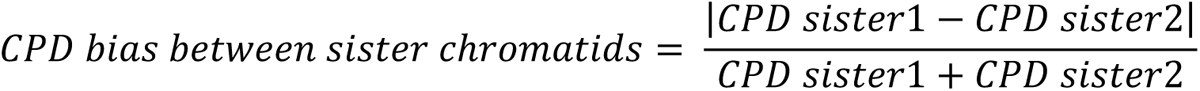

The maximum CPD bias was calculated by sorting together the chromatids bearing the highest CPD levels from each pair of damaged chromosomes (group 1) versus the chromatids bearing the lowest CPD levels (group 2) in each metaphase nucleus. The following formula was used to calculate the maximum CPD bias for each metaphase nucleus and converted into percent:

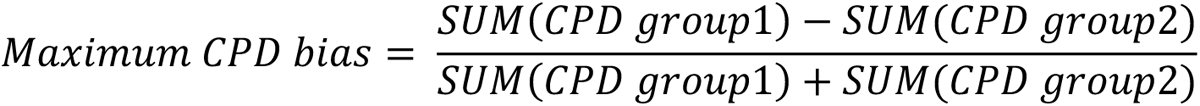

To simulate a random distribution of UV-damaged sister chromatids, we used the “RAND” function in excel to randomly assign each sister chromatid to group 1 or 2. This step was repeated for each damaged chromosome. The CPD bias expected from the simulated random segregation of sister chromatids was calculated as above.

To calculate the proportion of damaged telomeres in the two G1 daughter cells, TRF2 foci were identified on Image J with the function “Find Maxima”. The number of TRF2 foci colocalizing with UV damaged spots (CPD) was measured in each daughter cell and normalized to the total number of TRF2 foci in the two daughter cells and converted into percent.

The coefficient of variation of DAPI staining (DAPI CV) corresponds to the standard deviation divided by the mean intensity of DAPI staining which reflects the difference in intensity between 2 neighboring pixels i.e. the spatial heterogeneity of the DAPI signal. Therefore, condensed chromosome regions have a higher DAPI CV than decondensed chromosome regions.

Image J macros developed for the study are available upon request.

### Statistical analysis

Statistical tests were performed using GraphPad Prism. Loss or enrichment of histone modifications or proteins of interest at UV sites relative to the whole nucleus was compared to a theoretical mean of 1 by one-sample t-test (two-sided). p-values for mean comparisons between two groups were calculated with a two-sided Student’s t-test with Welch’s correction. p-values for mean comparisons between more than two groups were calculated with a one-way ANOVA (Gaussian distribution and assuming equal SD). The Dunnett’s correction was applied when samples were only compared with the control condition. Spearman correlation was applied to compare datasets. ns: non-significant, *: p < 0.05, **: p < 0.01, ***: p < 0.001, ****: p < 0.0001. Confidence interval of all statistical tests: 95%.

## AUTHOR CONTRIBUTIONS

J.F. and J.D. designed and performed experiments with technical assistance from O.C., and analysed the data. J.F. and S.E.P. wrote the manuscript. M. K-C performed experiments with the ATR inhibitor. A.K., J.H and S.R. tested the antibodies with peptide arrays. S.E.P. supervised the project. All authors edited and approved the final version of the manuscript.

## ACKNOWLEDGMENTS

We thank members of our laboratory for stimulating discussions and Slimane Ait-Si-Ali and Pierre-Antoine Defossez for critical reading of the manuscript. We thank H. Kimura and I. Matic for sharing antibodies and V. Mezger for sharing the iPS(IMR90)-4 cell line and antibodies. We acknowledge the help of the ImagoSeine facility (Institut Jacques Monod, France BioImaging) and of the imaging platform of the Epigenetics and Cell Fate Centre for confocal and epifluorescence microscopy. This work was supported by the European Research Council (starting grant ERC-2013-StG-336427 “EpIn” and consolidator grant ERC-2018-CoG-818625 “REMIND”), the French National Research Agency (ANR-18-CE12-0017-01 “HEROD”), the “Who am I?” laboratory of excellence (ANR-11-LABX-0071) funded by the French Government through its “Investments for the Future” program (ANR-11-IDEX-0005-01), and France-BioImaging (ANR-10-INBS-04). S.E.P. is an EMBO Young Investigator. J.F and J.D. received PhD fellowships from University Paris Diderot, Fondation ARC and EUR G.E.N.E (ANR-17-EURE-0013). S.B.R. acknowledges support from the National Institutes of Health (R35GM152184).

## COMPETING INTERESTS STATEMENT

The authors declare no competing interests.

## DATA AVAILABILITY

All data generated during this study are included in this article and its supplementary information files.

## Notes

### Competing Interest Statement

The authors have declared no competing interest.

### Summary of Updates

As main additions to the manuscript, we now provide a technically robust demonstration of non-random damage inheritance; we discard alternative hypotheses that could explain the asymmetric damage distribution; we explore the functional consequences of asymmetric damage segregation on DNA damage signaling and on stemness marker expression in daughter cells.

