## Supplementary figures and images for "Mitotic chromatin marking governs asymmetric segregation of DNA damage"

### Supplementary Figures

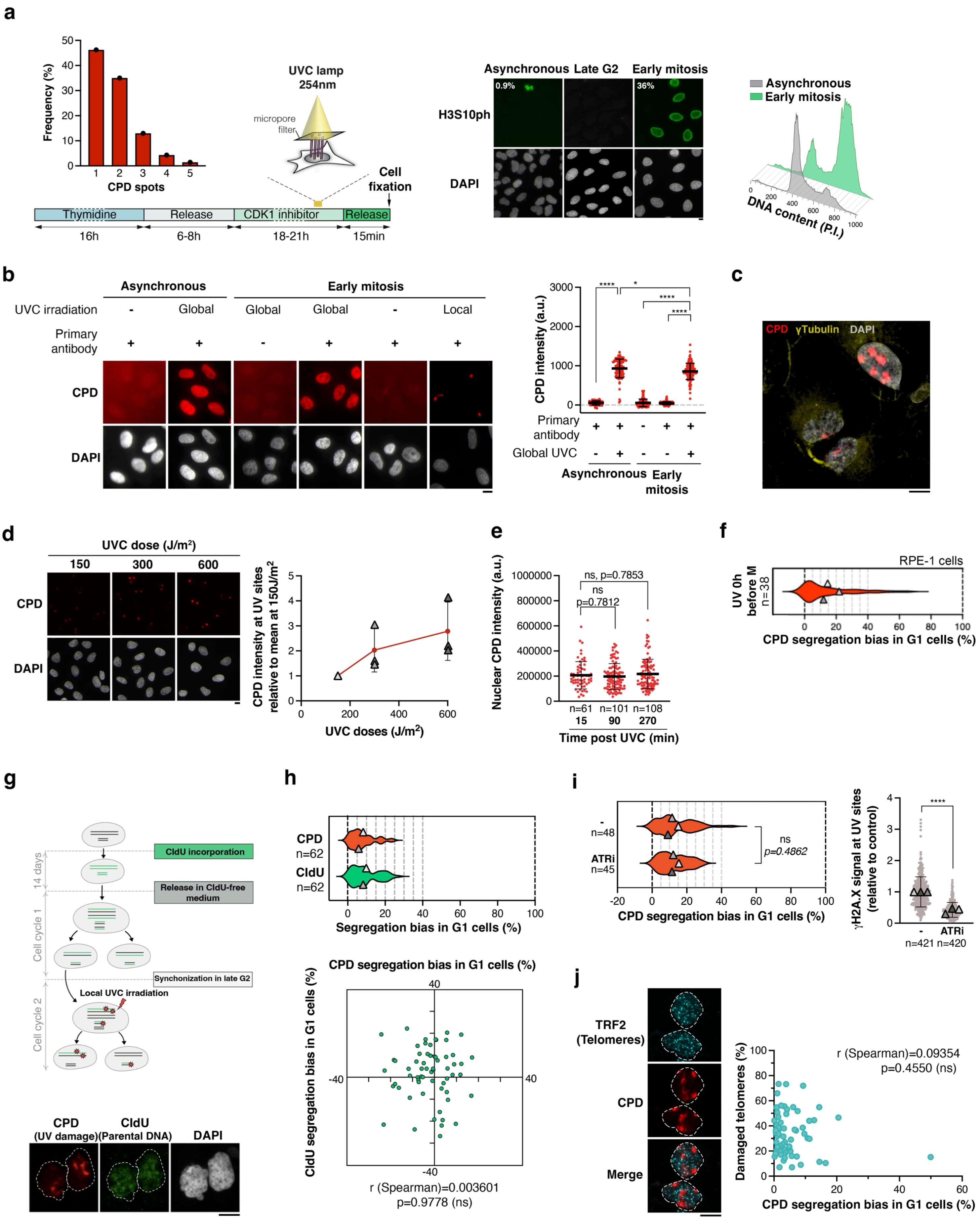

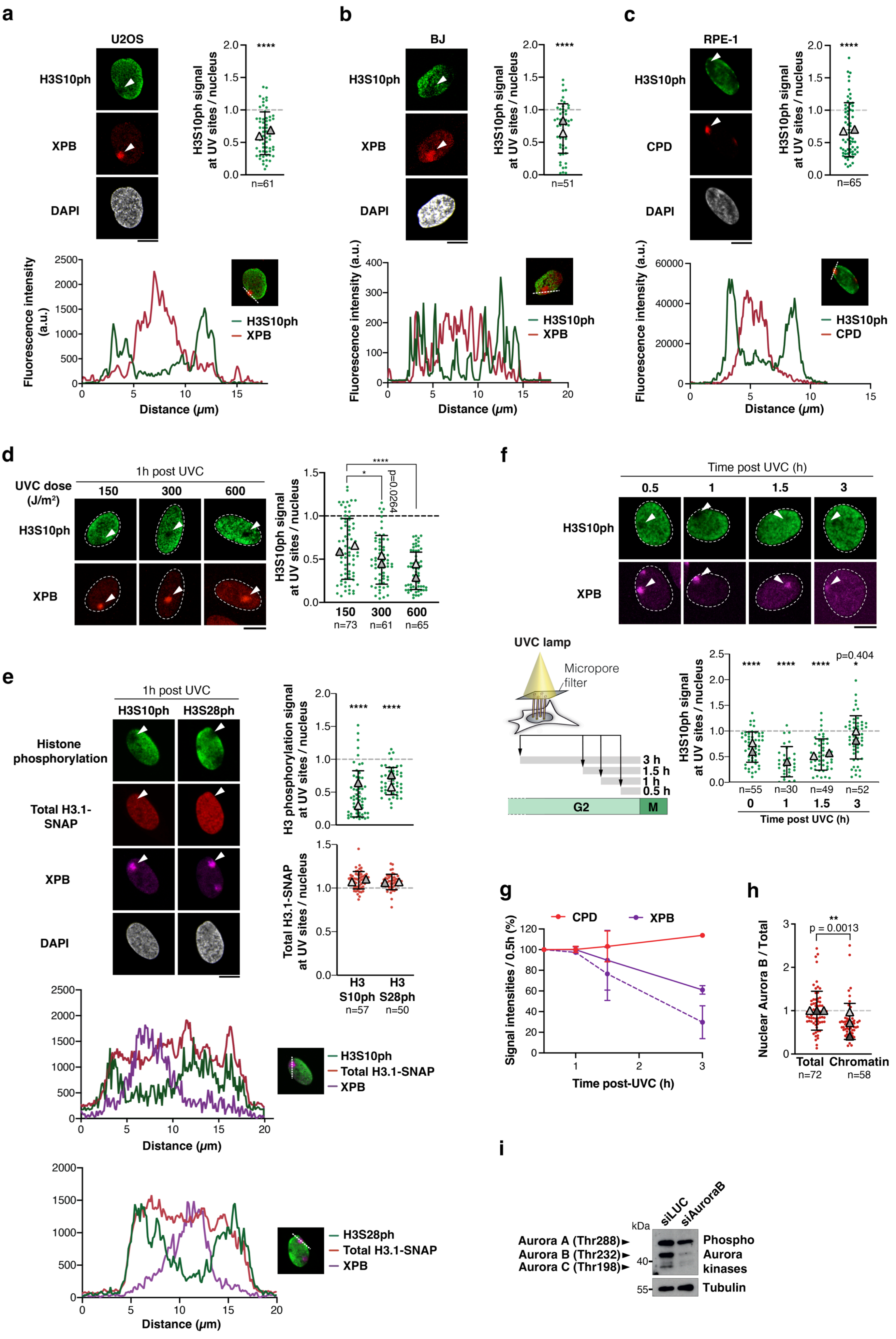

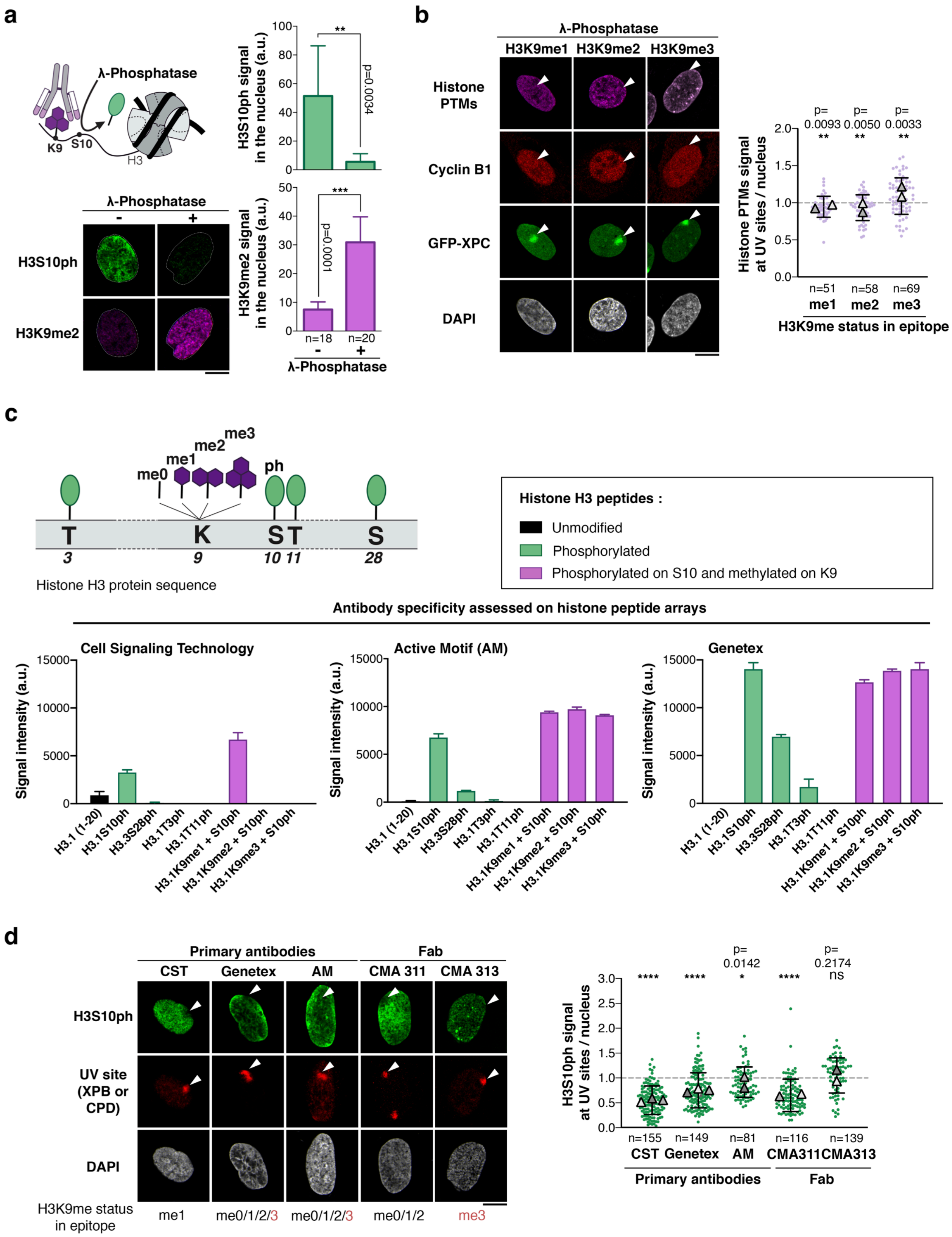

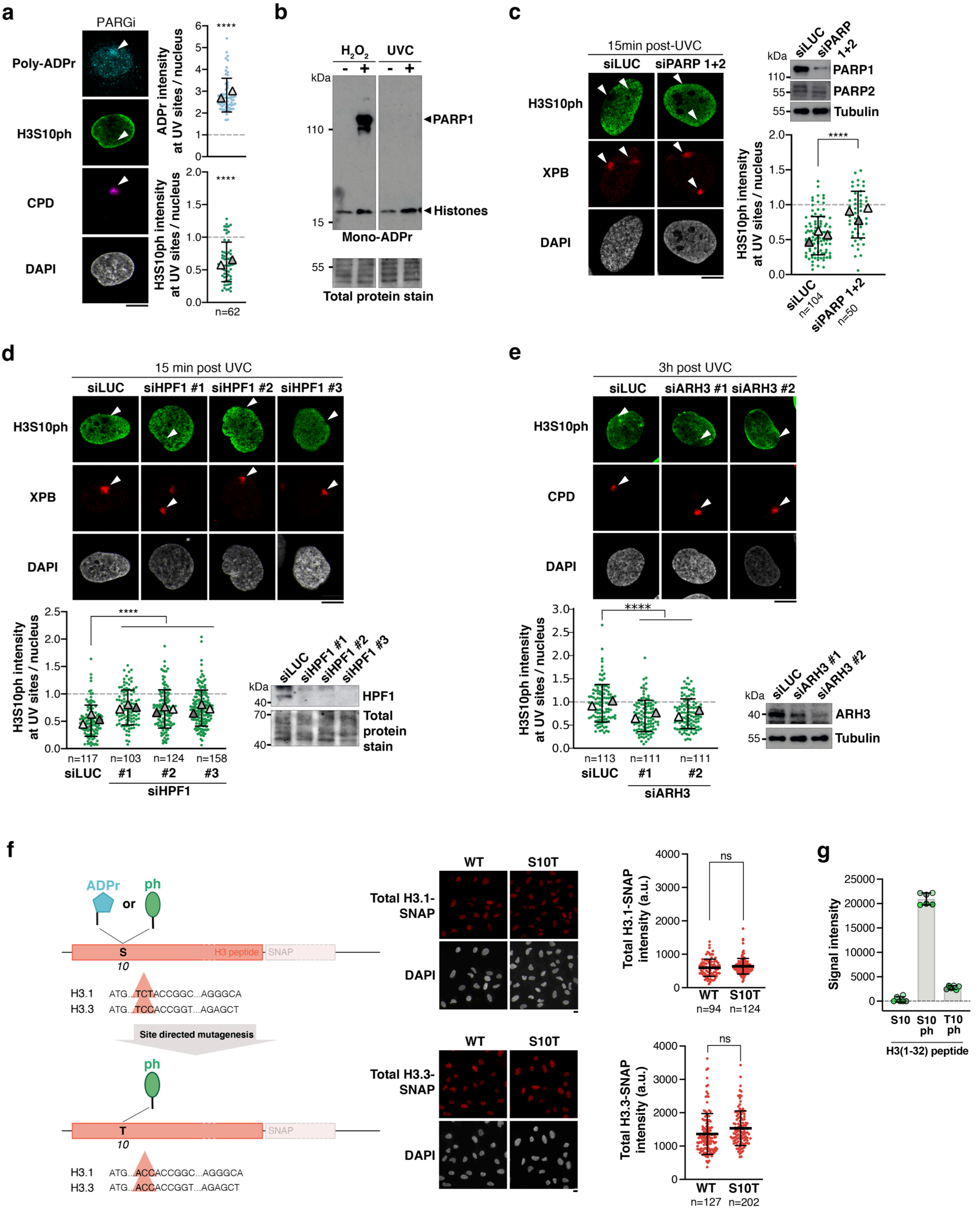

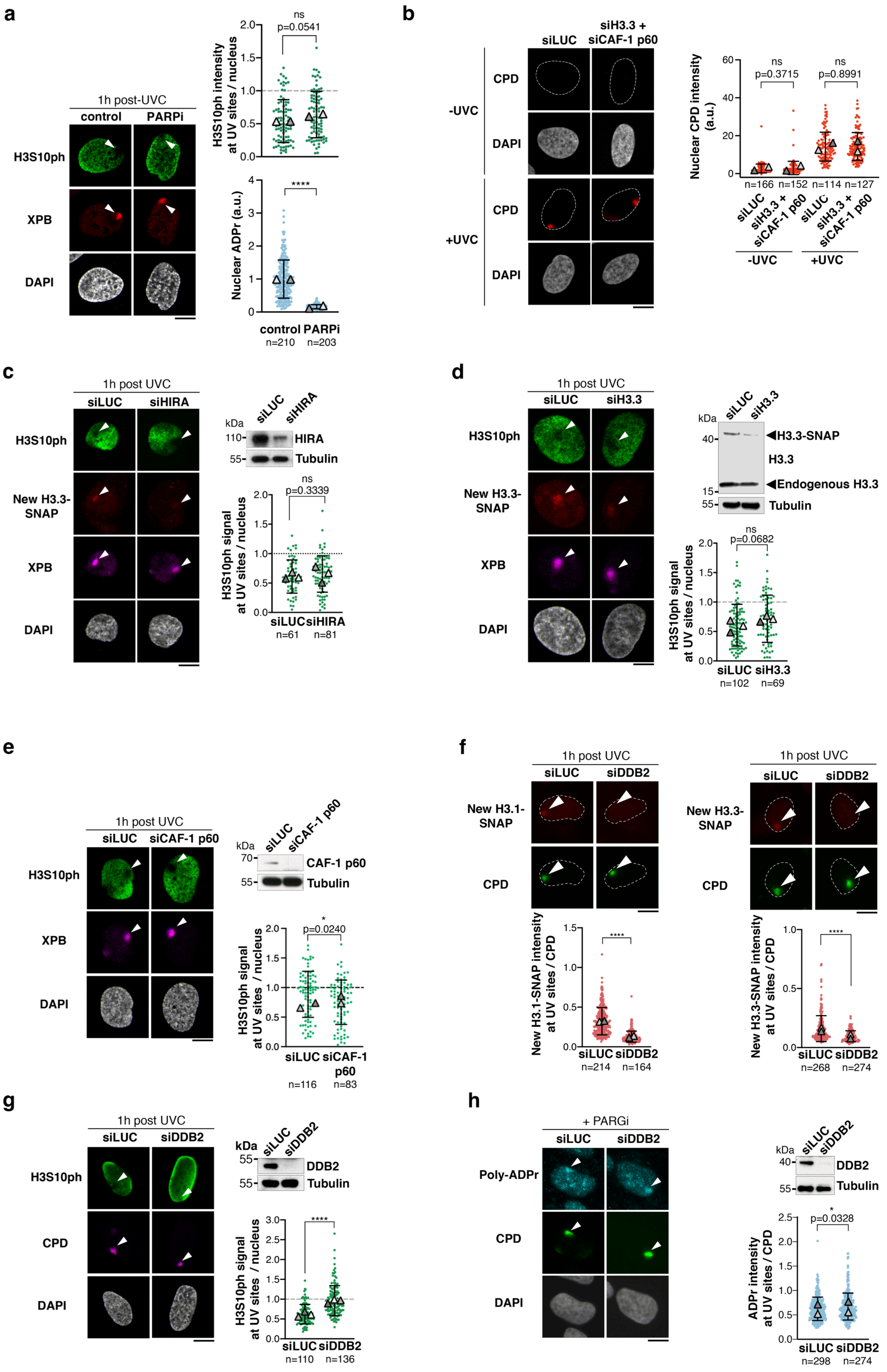

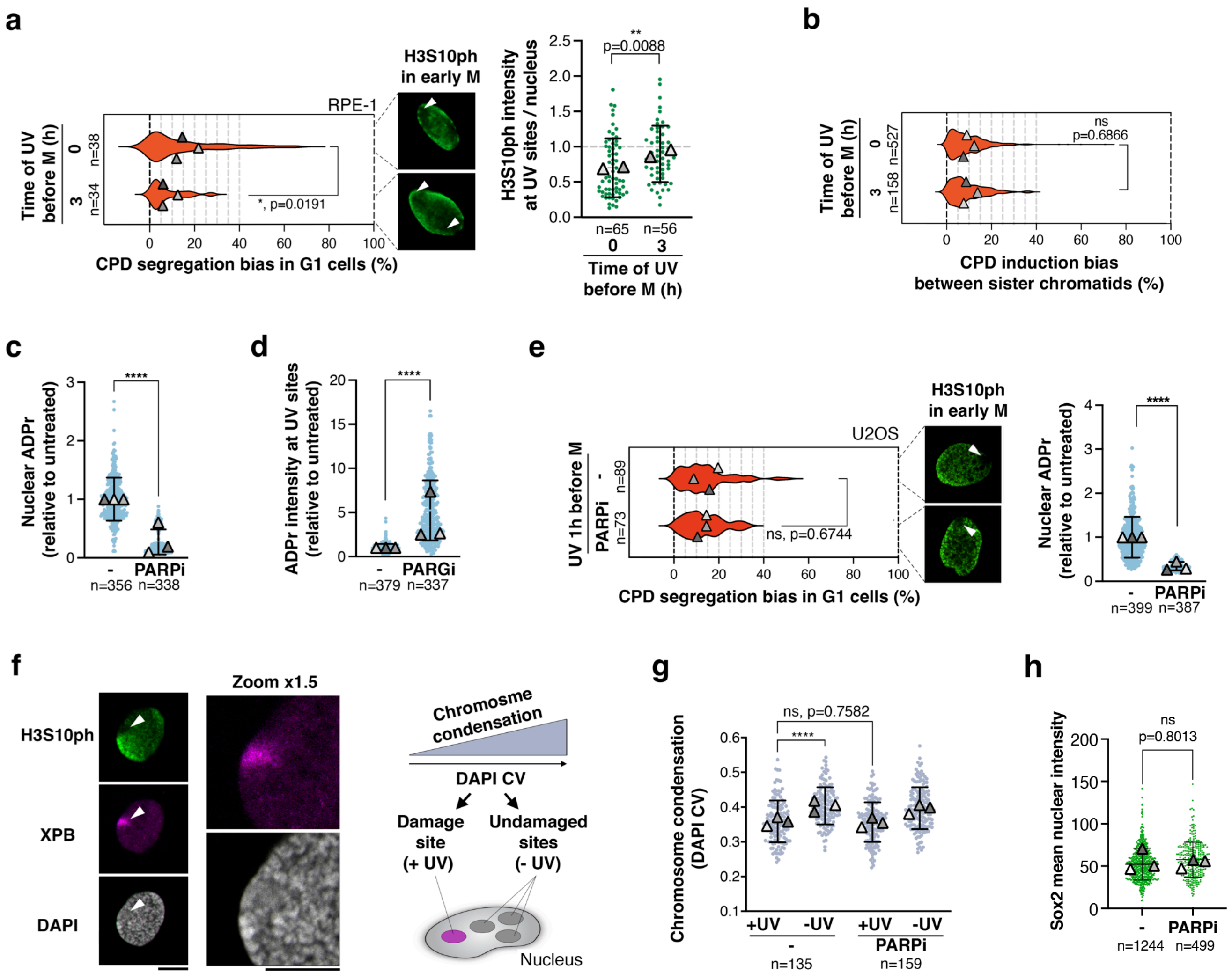
